# Structural and functional damage to neuronal nuclei caused by extracellular tau oligomers

**DOI:** 10.1101/2023.05.08.539873

**Authors:** Xuehan Sun, Guillermo Eastman, Yu Shi, Subhi Saibaba, Ana K. Oliveira, John R. Lukens, Andrés Norambuena, James W. Mandell, George S. Bloom

## Abstract

**INTRODUCTION:** Neuronal nuclei are normally smoothly surfaced. In Alzheimer’s disease (AD) and other tauopathies, though, they often develop invaginations. We investigated mechanisms and functional consequences of neuronal nuclear invagination in tauopathies.

**METHODS:** Nuclear invagination was assayed by immunofluorescence in brain, and in cultured neurons before and after extracellular tau oligomers (xcTauO) exposure. Nucleocytoplasmic transport was assayed in cultured neurons. Gene expression was investigated using nanoString nCounter technology and qRT-PCR.

**RESULTS:** Invaginated nuclei were twice as abundant in human AD as in cognitively normal adults, and were increased in mouse neurodegeneration models. In cultured neurons, nuclear invagination was induced by xcTauOs by an intracellular tau-dependent mechanism. xcTauOs impaired nucleocytoplasmic transport, increased histone H3 trimethylation at lysine 9 and altered gene expression, especially by increasing tau mRNA.

**DISCUSSION:** xcTauOs may be a primary cause of nuclear invagination *in vivo*, and by extension, impair nucleocytoplasmic transport and induce pathogenic gene expression changes.

## 1 INTRODUCTION

Oligomeric and filamentous tau are key pathogenic factors in tauopathies, such as Alzheimer’s Disease (AD), frontotemporal dementia (FTD) and progressive supranuclear palsy (PSP). Tauopathy spreads from neuron to neuron by cycles of tau aggregate release into the extracellular space and subsequent neuronal uptake [1–8]. While the mechanisms of this prion-like spread of pathogenic tau has been extensively studied, the cell biological responses of neurons to aggregated tau uptake has attracted much less attention.

The nuclear lamina, a meshwork of intermediate filaments formed by lamins A, B1, B2 and C, that underlies the inner nuclear membrane, plays critical roles in maintenance of normal nuclear structure and function. Deformation of the nuclear membrane, including nuclear lamina invagination, has been implicated in multiple neurodegenerative diseases, including AD, FTD, and Parkinson’s disease, and can be provoked by dysfunctional tau [9–11]. For example, expression of pathogenically mutated human tau in *Drosophila* neurons caused nuclear lamina invagination [11], and intracellular tau oligomerization led to disassembly of lamin B2, a major component of the nuclear lamina [12].

With this background in mind, we investigated whether extracellular tau oligomers (xcTauOs) affect neuronal nuclei. We present the novel findings that xcTauOs rapidly cause striking nuclear invagination, and that it occurs by a mechanism that depends on intracellular tau, and is associated with altered nucleocytoplasmic transport, chromatin structure and gene transcription. Altogether, our results implicate xcTauOs as seminal factors in the conversion of healthy neurons into diseased neurons in AD and other tauopathies.

## 2 METHODS

### 2.1 Human brain tissue

Paraffin-embedded, 5-6 μm thick human cortical brain autopsy sections were obtained from the archives of the University of Virginia Department of Pathology. Frozen human cortical brain autopsy samples were provided by Dr. Heather A. Ferris of the University of Virginia School of Medicine. Pertinent information about each donor is shown in Supplementary Table 1. Institutional approval for use of archival autopsy tissue was obtained from the University of Virginia Biorepository and Tissue Research Facility.

### 2.2 Mouse brain tissue

CVN mice were originally obtained from Drs. Michael Vitek and Carol Colton of Duke University, and were maintained as a breeding colony. PS19 mouse brain sections were provided by Dr. John Lukens of the University of Virginia Department of Neuroscience. Animals were maintained, bred, and euthanized in compliance with all policies of the Animal Care and Use Committee of the University of Virginia. 50 μm floating sections of brain tissue were cut following trans-cardiac perfusion of 4% paraformaldehyde in PBS of mice that had been deeply anesthetized intraperitoneally with ketamine/xylazine (280/80 mg/kg).

hTau mice were originally obtained from the Jackson Laboratory and maintained as a breeding colony. Mice were euthanized by carbon dioxide inhalation. The brains were then harvested and saved at −80°C for future use. For all experiments, only male mice were used.

### 2.3 Cultured mouse neurons

Brain cortices were collected from E17/18 wild type (WT) C57/Bl6 or tau knockout mice [13] and processed as described earlier [14]. Briefly, brain tissue was cut into small pieces in ice-cold Hank’s balanced salt solution, and then was digested at 37°C with 0.25% trypsin (Gibco, 15090-046) and 500 units DNase (Worthington, LK003170) for 20 minutes. Digestion was stopped by adding equal volume of fetal bovine serum. The resulting single cell suspension was washed three time with 5 ml of warm Neurobasal Plus medium (Gibco, A3582901) containing B27 Plus supplement (Gibco, A3582801), 2mM GlutaMAX (Gibco, 35050061), 6mg/ml glucose (Sigma-Aldrich, 16301) and 10 µg/ml gentamicin (Gibco, 15750078), mechanically dissociated with a fire polished Pasteur pipet and diluted into supplemented Neurobasal Plus medium. 65000 cells/cm^2^ were plated on 50 μg/ml poly-D-Lysine (Sigma-Aldrich, P0899) coated plates. Neurons were maintained in a tri-gas incubator in an atmosphere of 5% each of O_2_ and CO_2_. 50% media changes were done every 3-4 days until day 12 or 13 when experimental treatments began.

### 2.4 Recombinant tau

All 6 human tau isoforms with a C-terminal his-tag were produced by expression in BL21(DE3) *E. coli* cells. Protein expression was induced with 0.5 mM Isopropyl β-D-1-thiogalactopyranoside (Sigma-Aldrich, I6758) for 2 hours at 37°C. Cells were pelleted by centrifugation, resuspended and sonicated with Misonix Sonicator 3000 20 times for 30 seconds each at 60% power. Next, the *E. coli* lysate was centrifuged in Sorvall RC6 Plus 6 centrifuge at 10,900 rpm for 25min with an SLA 1500 rotor. Tau was then purified batchwise from the supernatant using TALON Metal Affinity Resin (TaKaRa, 635502) according to the vendor’s instructions. Finally, each purified tau isoform was concentrated, and buffer exchanged into 10mM HEPES, pH 7.6, using Amicon Ultra-4 Centrifugal Filters (Millipore, UFC801024).

### 2.5 Tau oligomers

Oligomeric tau was prepared as described earlier [15]. Each purified tau isoform was adjusted to 8 μM in the oligomerization buffer: 10 mM HEPES (pH 7.6), 100mM NaCl, 0.1 mM EDTA and 5mM DTT. The protein was then allowed to oligomerize in the presence of 300μM Arachidonic acid (Cayman Chemicals, 90010) for 18 hours at room temperature in the dark. Oligomerization was verified on western blots. Primary mouse neurons at 12-13 days *in vitro* were exposed to oligomeric or monomeric tau at 30-500 nM total tau, or with vehicle for the indicated time periods.

### 2.6 Lentiviruses

Human 2N4R tau cDNA was amplified by PCR and transferred to the FSW lentiviral vector between the BamHI and HpaI sites. Primers was described earlier [16]. The expression plasmids encoding tau, and the packaging vectors, (pSPAX2 and pMD2.G; (Addgene plasmids 12260 and 12259, respectively) were transfected using Lipofectamine 3000 (ThermoFisher, L3000001) into HEK293T cells grown in 15 cm Petri dishes to ∼80% confluence in Dulbecco’s Modified Eagle’s medium (Gibco 11965-092) supplemented with 10% fetal bovine serum (VWR, 89510-186). Each transfection employed 15 µg total DNA at a 50%/37.5%/12.5% ratio of expression vector/pSPAX2/pMD2.G. Lentivirus-conditioned medium was collected 24 and 48 hours after the start of transfection. Lentiviral particles were concentrated in a Beckman Coulter Optima LE-80K ultracentrifuge for 2 hours at 23,000 rpm at 4°C in an SW28 rotor, resuspended in 400 µl Neurobasal medium and stored at −80°C in small aliquots. Cultured neurons were transduced in Neurobasal/B27 medium and incubated for 48-72 hours before assays were performed.

### 2.7 Brain homogenates

Cerebral hemispheres were dissected from freshly euthanized mice, suspended in 3 volumes of Neurobasal plus medium at 4° C, and homogenized on ice using 25 pulses of 30 seconds each of a probe sonicator (Misonix Sonicator 3000) at 30% power. Lysates were centrifuged at 21,000 g for 15min with a TLA 120.2 rotor (Beckman Optima TLX Ultracentrifuge) to remove cellular debris and large, insoluble material. Supernatants were then passed through a 0.22 um filter, aliquoted and stored at −80 °C until further use. Total protein concentration was determined by Pierce BCA Protein Assay Kit (Thermo Scientific, 23225).

### 2.8 Quantitative immunofluorescence

All micrographs were acquired using an inverted Nikon Eclipse Ti microscope equipped with a Yokogawa CSU-X1 spinning disk confocal head with 405 nm, 488 nm, 561 nm and 640 nm lasers, and 10X and 20X dry, and 40X and 60X oil immersion Nikon Plan Apo objectives. All brain and cultured neuron immunofluorescence micrographs are shown as maximum projections of Z-stacks produced using the <u>Fiji</u> derivative of <u>ImageJ.</u> Primary and secondary antibodies are listed in Supplementary Table 2.

Primary mouse neurons growing on coverslips were rinsed once with PBS, and with one exception, were fixed and permeabilized in methanol for 5 minutes at −20° C. The exception was that cells stained with anti-Ran were fixed with 2% paraformaldehyde for 10 minutes at room temperature and permeabilized with 0.2% Triton X-100 for 10 minutes. After washing three times with PBS, cultured neurons were blocked in Intercept (PBS) Blocking Buffer (LI-COR, 927-70001) /0.1% Tween 20 for 1 hour and incubated with the indicated primary antibodies for 30 minutes, followed by the indicated secondary antibodies for 30 minutes. All antibodies were diluted in Intercept (PBS) Blocking Buffer/0.1% Tween20. After each antibody incubation step, the cells were rinsed 3 times for 5 minutes each with PBS. In the last wash, the coverslips were incubated with 5ug/ml DAPI (Sigma-Aldrich, D9542). The coverslips were then mounted onto slides using Fluoromount-G (Southern Biotech, 0100-01). To quantify nuclear invagination in primary neurons, lamin B1-positive pixels were assigned as either nuclear boundary or invaginated using a Fiji thresholding algorithm. The ratio between the area of invaginated lamin B1 over that of the total laminB1 staining was reported (Figure 2A). At least 100 neurons per condition were evaluated fort each experiment.

Human paraffin embedded brain sections were first deparaffinized and rehydrated by sequential incubation in Xylenes (2 x 5 minutes), 1:1 xylenes:100% ethanol (3 minutes), 100% ethanol (3 minutes), 95% ethanol (3 minutes), 70% ethanol (3 minutes), 50% ethanol (3 minutes), and distilled water (3 minutes). Antigen retrieval was achieved by microwaving the sections in citrate buffer pH 6.0 (Vector Laboratories, H-3300) for 15 minutes. After cooling to room temperature, sections were rinsed in PBS and blocked in PBS/5% BSA/0.1% Triton X-100 for 1 hour at room temperature and incubated with the indicated primary antibodies diluted in PBS/5% BSA overnight at 4° C. Sections were then washed 4 times for 5 minutes each in PBS and incubated with the indicated secondary antibodies for 2 hours at room temperature. After washing 4 times for 5 minutes each in PBS, sections were incubated with DAPI for 10 minutes and then with autofluorescence eliminator (Millipore, 2160) for 5 minutes, followed by 3 washes with ethanol. Finally, the tissue sections were mounted under #1 thickness coverslips using Fluoromount-G. Approximately 50 MAP2-positive nuclei were evaluated in each case. A nucleus was considered invaginated if the total length of its invaginations were at least 10% of the longest axis of the nucleus.

Free floating, 50 µm mouse brain sections were rinsed in PBS for 5 minutes and blocked with PBS/5% normal goat serum (Southern Biotech, 0060-01) for 2 hours at room temperature. Sections were then incubated with the indicated primary antibodies diluted into PBS/2% normal goat serum/0.05% Tween 20 overnight at 4° C. Sections were then washed with PBS/0.05% Tween 20 3 times for 10 minutes each and incubated with the indicated secondary antibodies for 2 hours at room temperature. After three washes with PBS, the sections were incubated with DAPI for 10 minutes and autofluorescence eliminator for 5 minutes, followed with 3 washes with ethanol. After the final wash, the sections were rinsed with PBS and mounted between #1 thickness coverslips and glass slides using Fluoromount-G. To ensure thorough neuroanatomical coverage, cortical sections from anterior, middle and posterior regions were imaged and about 300 neurons from each section were evaluated. A nucleus was considered invaginated if the total length of its invaginations were at least 10% of the longest axis of the nucleus.

### 2.9 Protein electrophoresis and western blots

For cultured neurons, total protein was extracted using RIPA buffer (Thermo Scientific, 89900) supplemented with protease inhibitors (Thermo Scientific, 78430) and phosphatase inhibitors (Thermo Scientific, 78420). Protein samples were mixed with 1X NuPage LDS Sample Buffer (Invitrogen, NP0007) and 1x sample reducing agent (Invitrogen, NP0004). Samples were heated at 70°C for 10 minutes and separated on 4-12% gradient Bis-Tris SDSPAGE gels (Invitrogen, NP0323) and transferred to 0.2μm nitrocellulose membrane (Bio-Rad Laboratories, 1620112) for 1 hour at 100V.

For human brain tissue, 50-100 mg of tissue was blended with one scoop of zirconium oxide beads (Next Advance, ZOB05) in 300ul buffer containing 65mM Tris-HCl, pH 6.8, 2.1% SDS, 5% β-mercaptoethanol and 0.1% Triton X-100 for 10 minutes. The lysate was then sonicated for 5 minutes and centrifuged at 14,000 g for 15 minutes with an accuSpin Micro 17R Centrifuge (Fisher Scientific) at 4°C. Samples were diluted 5-fold into 1x Laemmli sample buffer (Bio-Rad Laboratories, 1610747) with 5% β-mercaptoethanol. The samples were then heated at 95°C for 10 minutes, separated on 4-12% gradient Bis-Tris SDSPAGE gels (Invitrogen, NP0323), and transferred to 0.2μm nitrocellulose membrane (Bio-Rad Laboratories, 1620112) for 8 hours at 25V.

Primary and secondary antibodies for western blotting are listed in Supplementary Table 2. All antibodies were diluted in Intercept (TBS) Blocking Buffer/0.1% Tween-20. Following protein transfer, nitrocellulose membranes were blocked with Intercept (TBS) Blocking Buffer (LI-COR, 927-60001) for 1 hour at room temperature and incubated with the indicated primary antibodies overnight at 4° C. Membranes were then washed 3 times for 5 minutes each in TBST (TBS/0.1% Tween 20) before being incubated with the indicated secondary antibodies at room temperature for 1 hour. Finally, the membranes were washed 3 times in TBST, and were imaged using a ChemiDoc MP imager and analyzed using Image Lab software (Bio-Rad).

### 2.10 Dextran Exclusion Assay

A previously described protocol [17] was adapted to primary WT mouse neurons that grew on glass coverslips and were exposed to xcTauOs (250nM total tau) or vehicle for 24 hours, and then washed once with prewarmed PBS and ice-cold PBS, respectively. Next, neurons were incubated in permeabilization buffer (20 mM HEPES pH 7.5, 110mM KOAc, 5 mM MgCl_2_, and 0.25 M sucrose) for 5 minutes on ice, followed by a 7-minute incubation with permeabilization buffer containing 20 µg/mL of digitonin (Sigma-Aldrich, CHR103). Neurons were then washed 4 times with diffusion assay solution (20 mM HEPES pH 7.5, 110 mM KOAc, 5 mM NaCl, 2 mM MgCl_2_, 0.25 M sucrose) on ice and incubated with 1 mg/ml of fluorescein-dextran (Invitrogen, D1823) for 10 minutes at room temperature. Hoechst 33342 was used as the nuclear counterstain (NucBlue Live ReadyProbe Reagent, Invitrogen, R37605). The glass coverslips were sealed onto slides with clear nail polish and imaged using a 20X 0.4 NA Plan Fluorite objective on an EVOS M5000 cell imaging system (ThermoFisher Scientific). Images were quantitatively analyzed using Fiji software: regions of interest (ROIs) defining nuclei region were hand drawn based on Hoechst staining, and the mean pixel intensity of nucleus-associated fluorescein-dextran was standardized against the mean pixel intensity of the background.

### 2.11 Specificity validation of an antibody to histone H3 trimethylated at lysine 9 (H3K9Me3)

Specificity of anti-H3K9Me3 was tested against two types of substrates: recombinant histone H3 (New England BioLabs, M2507S), which lacked methylation, and native histones that contain the modification and were purified from HEK293T cells using a Histone Extraction Kit (Abcam ab113476). Protein concentration was determined using the Pierce BCA assay. Samples were diluted into 1X NuPage LDS Sample Buffer (Invitrogen, NP0007), supplemented 1x sample reducing agent (Invitrogen, NP0004). 2.5 µg of native mixed histones and 625 ng of pure recombinant histone H3 were loaded on the gel.

### 2.12 Gene expression analysis and qRT-PCR

The nanoString nCounter system was used for quantitatively analyzing gene expression using the Mouse Neuropathology panel of 760 mRNA probes. Total RNA was isolated from cultured mouse neurons treated with xcTauOs (250 nM total tau) or vehicle for 6 hours using the mirVana isolation kit (Invitrogen, AM1560). RNA integrity was evaluated by capillary electrophoresis using an Agilent Bioanalyzer. 100 ng of RNA with a RIN >7 was used per sample, and 3 biological replicates were used for each condition. Data analysis was done using the ROSALIND platform considering a p-value threshold of 0.05 for differential gene expression; batch effects were considered in the analysis.

Semi quantitative real time PCR was used to quantify *MAPT* mRNA level in cultured mouse neurons after xcTauO or vehicle treatment for 6 hours. RNA was isolated using Trizol (Invitrogen, 15596026) and *MAPT* mRNA levels were quantified using One-Step qRT-PCR (Invitrogen, 11746100) in an Applied Biosystem StepOne Plus Real-Time PCR instrument as follows: 3 minutes at 50° C; 5 minutes at 95° C; 40 cycles of 15 seconds at 95 °C, 30 seconds at 60° C and 1 minute at 40° C; and 72° C to 90° C melting analysis. In all cases, *Actb* and *Rpl19* were used as reference genes, and 4 independent biological replicates with 2 technical replicates each were quantified. Primer sequences were obtained from Origene (MP208179, MP200232 and MP212857). Primer efficiency was calculated and incorporated in the ΔΔCt method analysis [18].

### 2.13 Statistical analysis

Data are presented as mean values of the number of independently conducted experiments indicated in the legend of each figure. Error bars represent the standard error of the mean (SEM). Statistical analysis was performed using Prism 9 software (GraphPad). Statistical tests used for all figures are indicated in the corresponding legends.

## 3 RESULTS

### 3.1 Nuclear lamina invagination in human AD and transgenic mouse brains

We first evaluated the morphology of neuronal nuclei in cortices from AD and age-mated cognitively normal individuals (Supplementary Table 1 shows clinical characterizations). Invaginated neuronal nuclei were identified by surrounding MAP2-positive cytoplasm and deep infoldings of the nuclear lamina, as determined by anti-lamin B1 immunofluorescence. ∼70% of neuronal nuclei were invaginated in AD brains, whereas only ∼30% showed such deformation in cognitively normal brains (Figure 1A). Quantitative western blotting of human prefrontal cortex detected 2 lamin B1 bands (Figure 1B), whose apparent molecular weights correspond to isoforms reported in the UniProtKB database, and the lamin B1 level in AD tissue was ∼28% lower than in tissue from cognitively normal individuals.

**FIGURE 1.**
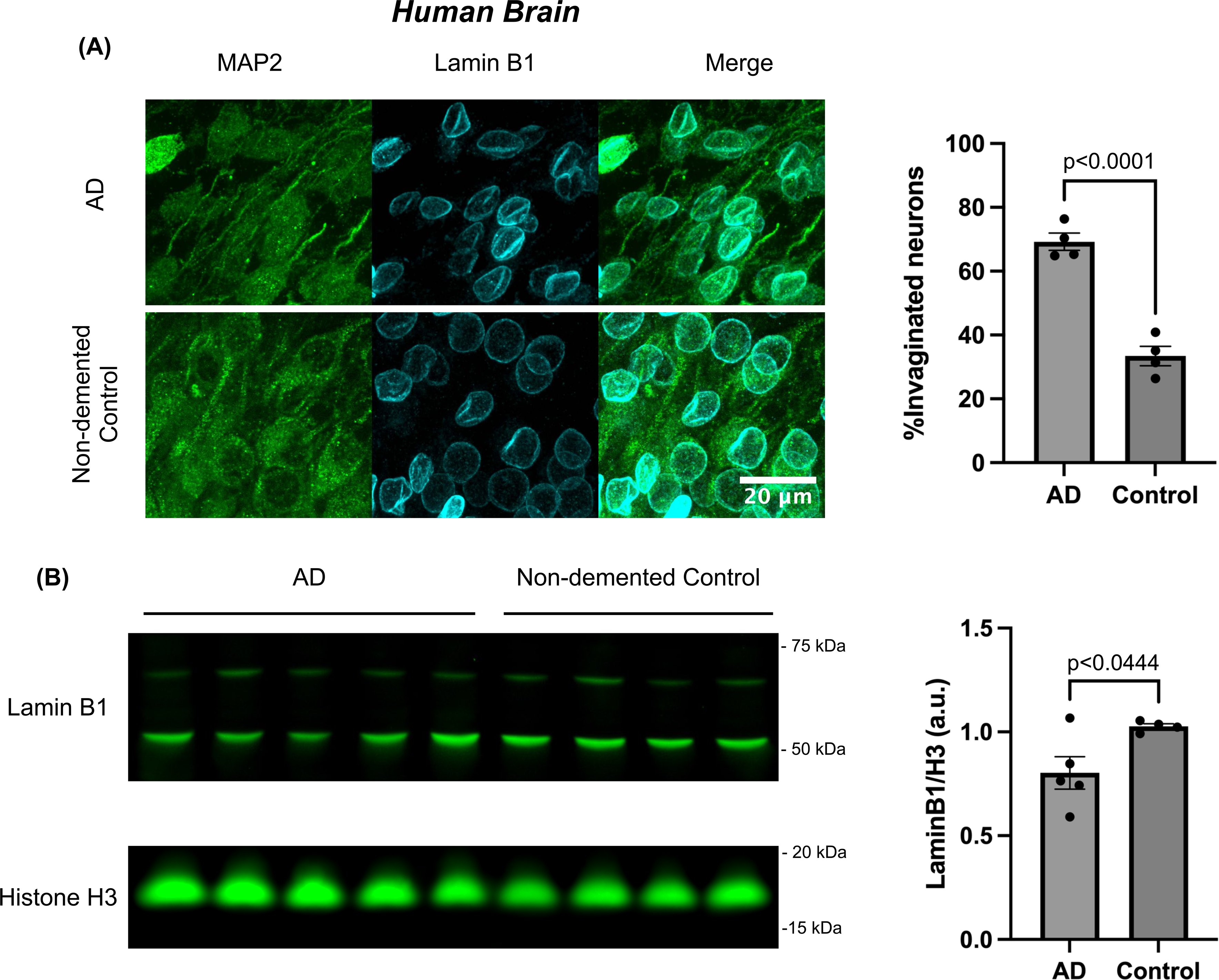
Nuclear lamina invaginations in human cortical AD neurons *in vivo*. (A) Immunofluorescence localization of the lamina protein, lamin B1, in MAP2-positive neurons, and quantitation of invaginated lamina. Quantitative data were obtained from 4 AD and 4 control cases (Supplementary Table 1), and for each case at least 50 neurons were scored. Significance was determined using unpaired t test, and error bars represent SEM. (B) Quantitative western blotting of lamin B1 in prefrontal cortices of human AD and non-demented control brains. For quantitation, lamin B1 signals were normalized against histone H3 for 5 AD and 4 normal cases (Supplementary Table 1). Significance was determined using unpaired t test, and error bars represent SEM.

We next studied nuclear morphology in 2 transgenic mouse lines: PS19, which overexpress human P301S tau, and accumulate tangles by 6 months and neuron loss within 9 months [19]; and CVN, which overexpress human APP with the Swedish, Dutch and Iowa mutations, are knocked out for nitric oxide synthase 2, and develop plaques, tangles and neuron loss within 12 months [20]. To ensure thorough neuroanatomical coverage, we sampled coronal sections from anterior, middle, and posterior brain regions for PS19 mice (Supplementary Figure 1), and lateral, middle, and medial sagittal sections for CVN mice (Supplementary Figure 2). Necab1, which is highly expressed only in layer IV neurons [21], was used to differentiate neurons in inner cortical layers to those in outer layers (Supplementary Figure 1A). Compared to age-matched WT mice, 6-month-old PS19 mice had higher levels of invaginated neuronal nuclei, especially in deep cortical layers. Elevated levels of such nuclei were also found in deep cortical layers of lateral brain regions in CVN mice (Supplementary Figure 2). Altogether, these findings indicate that invaginated neuronal nuclei are abundant *in vivo* in human AD brain, and in the brains of transgenic mouse models of AD and a pure tauopathy.

### 3.2 Human xcTauOs induce nuclear invagination and endogenous tau aggregates in cultured mouse neurons

Previous studies established that pathogenically mutated intracellular tau can provoke neuronal nuclear deformation [9, 11], but possible roles for xcTauOs [22] in this process have not been reported. We therefore investigated effects of xcTauOs on neuronal nuclei. All 6 human CNS tau isoforms were expressed in bacteria and purified, (Supplementary Figure 3), and oligomerized by addition of arachidonic acid (ARA; Figure 2B) [15]. The number of tau subunits in oligomers with apparent molecular weights of ∼110-130 kDa and ∼200 kDa seen on our western blots is unknown, but TauOs can migrate anomalously in SDSPAGE, as ∼180 kDa oligomers made from 2N4R tau by a similar method are actually dimers as determined by mass spectrometry [23]. We thus conclude that the TauOs used here represent low-*n* oligomers of undetermined specific size.

**FIGURE 2.**
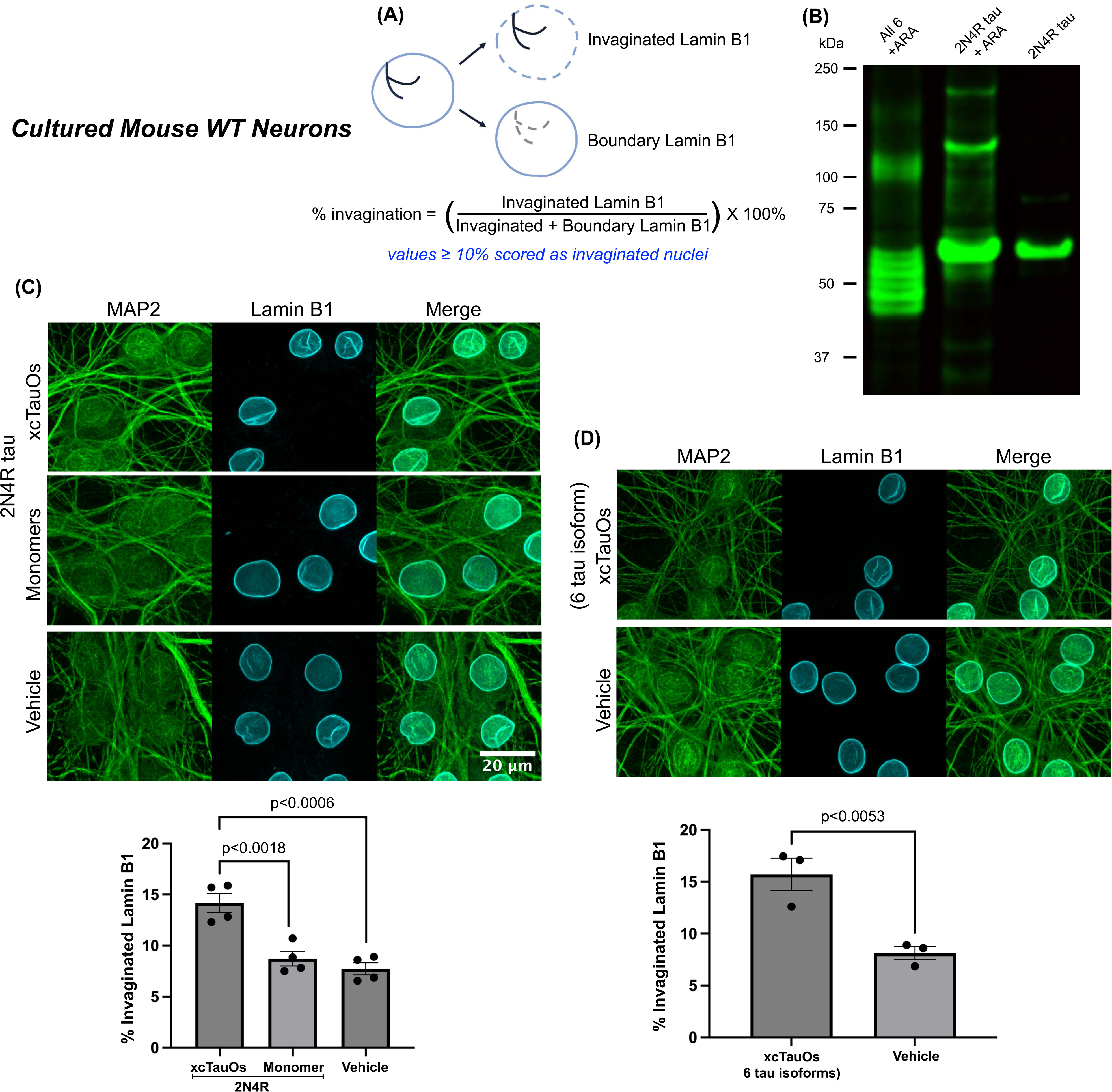
xcTauOs induce lamina invagination in cultured primary WT mouse cortical neurons. (A) Criteria for classifying nuclei as invaginated. For each nucleus, each lamin B1-positive pixel was assigned as either on the boundary or in an invagination. The ratio between the area of invaginated lamin B1 over that of the total laminB1 staining was reported.(B) Tau5 western blot of all 6 recombinant human CNS tau isoforms oligomerized with arachidonic acid (ARA), and recombinant human 2N4R tau (Supplementary Figure 3) with (oligomers) and without (monomers) ARA treatment. (C) Immunofluorescence localization of lamin B1 in (MAP2-positive) neurons treated for 1 hour with xcTauOs or tau monomers (250 nM total 2N4R tau in both cases), or vehicle. (D) Immunofluorescence localization of lamin B1 in (MAP2-positive) neurons treated for 1 hour with xcTauOs (250 nM total tau; an equimolar mix of all 6 human tau CNS isoforms), or vehicle. Significance was determined using Student’s t test; error bars represent SEM; n = 3 biological replicates, and at least 100 nuclei were scored per condition for C and D.

Nuclear invagination was induced in neurons exposed to xcTauOs made from 2N4R tau or an equimolar mixture of all 6 isoforms, but not to tau monomer or vehicle controls (Figure 2). Compared to controls, xcTauOs increased the extent of neuronal nuclear invagination by 63-95%. Dose-response and kinetic analyses established that xcTauOs made from 250nM total 2N4R tau caused maximum nuclear invagination within just 1 hour of treatment (Supplementary Figure 4). Because oligomers represented an average of ∼30% of the total tau, and may have been mainly dimers, the total oligomer concentration that was actually ∼30-40 nM.

To evaluate the *in vivo* relevance of xcTauOs assembled from recombinant tau, we also prepared and tested soluble brain extracts from 21-month-old hTau^+/-^ transgenic mice that express approximately equimolar levels of the 6 CNS tau isoforms encoded by human genomic tau DNA in the absence of full-length mouse tau [24]. Brain extracts from comparably aged littermates that were null for human tau (hTau^-/-^) were used as controls. Western blots revealed that all hTau^+/-^ brains contained detectable tau, but only some harbored TauOs, typically with a size range of ∼100-160 kDa (Figure 3A). hTau^+/-^ brain extracts containing TauOs plus tau monomers, but not monomers alone, potently induced neuronal nuclear invagination (Figure 3A). The results described so far in this section indicate that xcTauOs made from recombinant tau can reliably substitute for TauOs made *in vivo* in brain, and that xcTauOs made from recombinant 2N4R tau are as effective as those made from a cocktail of all 6 recombinant CNS tau isoforms. Accordingly, all other experiments described in this report relied on xcTauOs made from recombinant 2N4R human tau.

**FIGURE 3.**
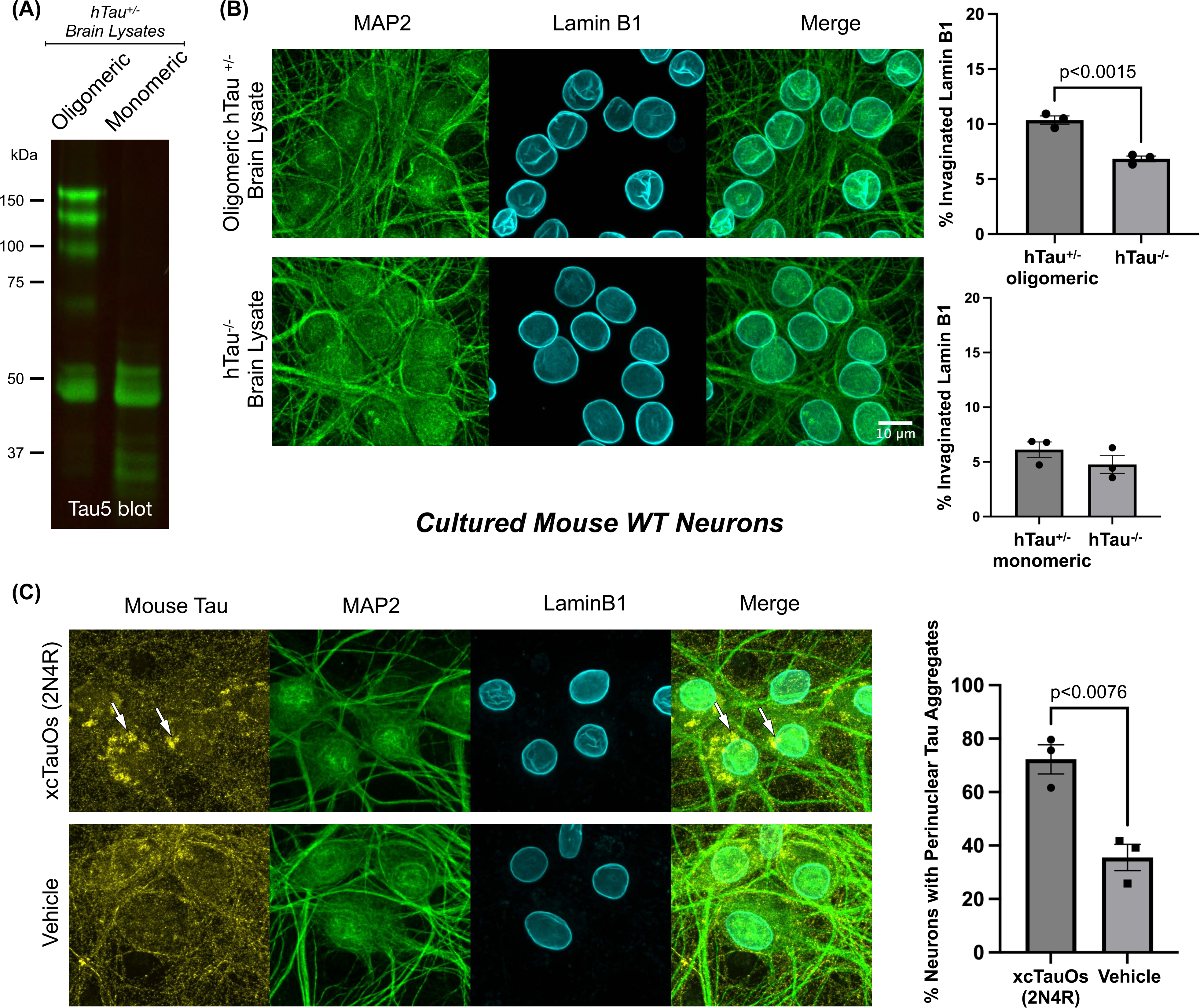
TauOs derived from hTau mouse brain induce lamina invagination and xcTauOs cause endogenous tau aggregation. (A) Tau5 western blot of hTau^+/-^ mouse brain extracts containing partially oligomeric or entirely monomeric tau. (B) Lamina invagination in primary mouse cortical neurons after a 4-hour exposure to hTau^+/-^ mouse brain extracts containing tau oligomers (diluted to 0.7% v/v). (C) A 1-hour treatment of primary mouse cortical neurons with xcTauOs made from recombinant human 2N4R tau (250 nM total tau) causes aggregation of endogenous, perinuclear mouse tau (white arrows). Significance was determined using Student’s t test; error bars represent SEM; n = 3 biological replicates for B and C.

In a prior study, we reported that xcTauOs made from recombinant human tau induce accumulation of intracellular tau aggregates in cultured mouse neurons, but we did not confirm the presence of endogenous mouse tau in those aggregates [25]. As shown in Figure 3B, conspicuous, perinuclear aggregates labeled with an antibody that recognizes mouse, but not human tau were abundant in xcTauO-treated cultured neurons.

### 3.3 The effect of xcTauOs on nuclear lamina architecture depends on intracellular tau

A recent report that intracellular TauOs causes translational stress [12] prompted us to test whether the mouse tau aggregates that formed in cultured mouse neurons exposed to human xcTauOs signal a requirement of intracellular tau for neuronal nuclear invagination induced by the xcTauOs. Accordingly, neurons derived from tau^-/-^ mice [13] were exposed to xcTauOs. As shown in Figure 4, neuronal nuclei in tau^-/-^ neurons did not invaginate after xcTauO exposure, unless human (2N4R) tau was expressed in the cells by lentiviral transduction. We therefore conclude that the effects of xcTauOs on nuclear morphology depend on intracellular tau.

**FIGURE 4.**
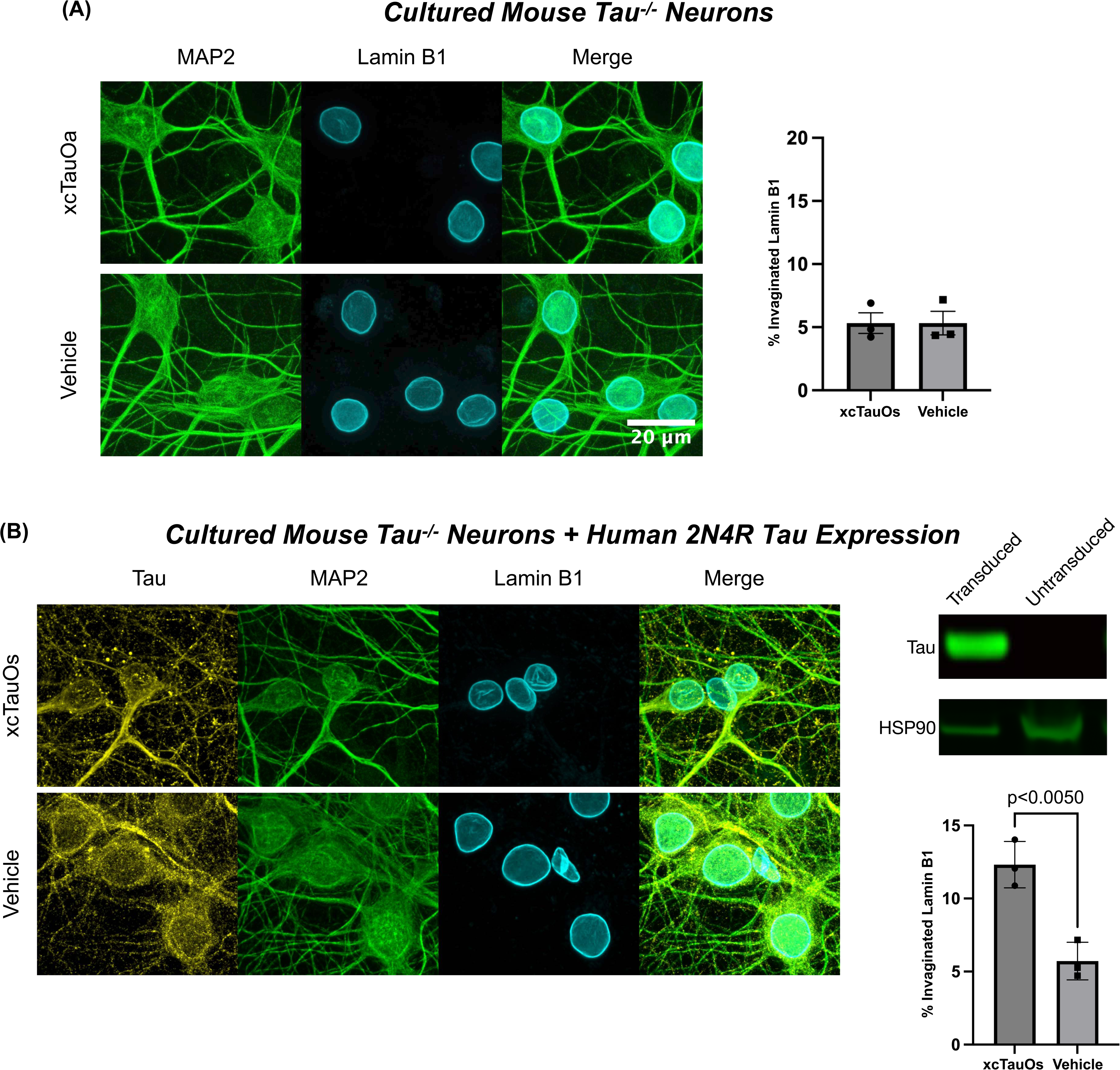
The xcTauO effect requires intracellular tau. (A) xcTauOs do not induce lamina invagination in primary cortical neurons derived from tau^-/-^ mice. Cultures were treated with xcTauOs (250 nM total tau) or vehicle for 1 hour. (B) Lentiviral expression of human 2N4R tau in tau^-/-^ neurons restores nuclear lamina sensitivity to a 1-hour exposure to xcTauOs (250 nM total tau). Significance was determined using Student’s t test; error bars represent SEM; n = 3 biological replicates for A and B.

### 3.4 xcTauOs disrupt nucleocytoplasmic transport in cultured neurons

The nuclear lamina provides a structural framework for nuclear pore complexes (NPCs), which regulate exchange of molecules larger than ∼40 kDa globular proteins between the nucleus and the cytoplasm. Nucleocytoplasmic transport is negatively impacted by mutant lamins [26, 27], which prompted us to assess whether it is also functionally compromised by xcTauOs. We first tested this possibility by a dextran exclusion assay, in which digitonin-permeabilized cultured mouse neurons were exposed to fluorescein-tagged 70 kDa dextran. In this assay, NPCs that are structurally damaged allow greater penetration of the dextran-fluorescein into the nucleus than undamaged NPCs (Figure 5A). We found that xcTauOs caused ∼36% more dextran-fluorescein to enter nuclei of neurons exposed to xcTauOs as compared to vehicle (Figure 5B).

**FIGURE 5.**
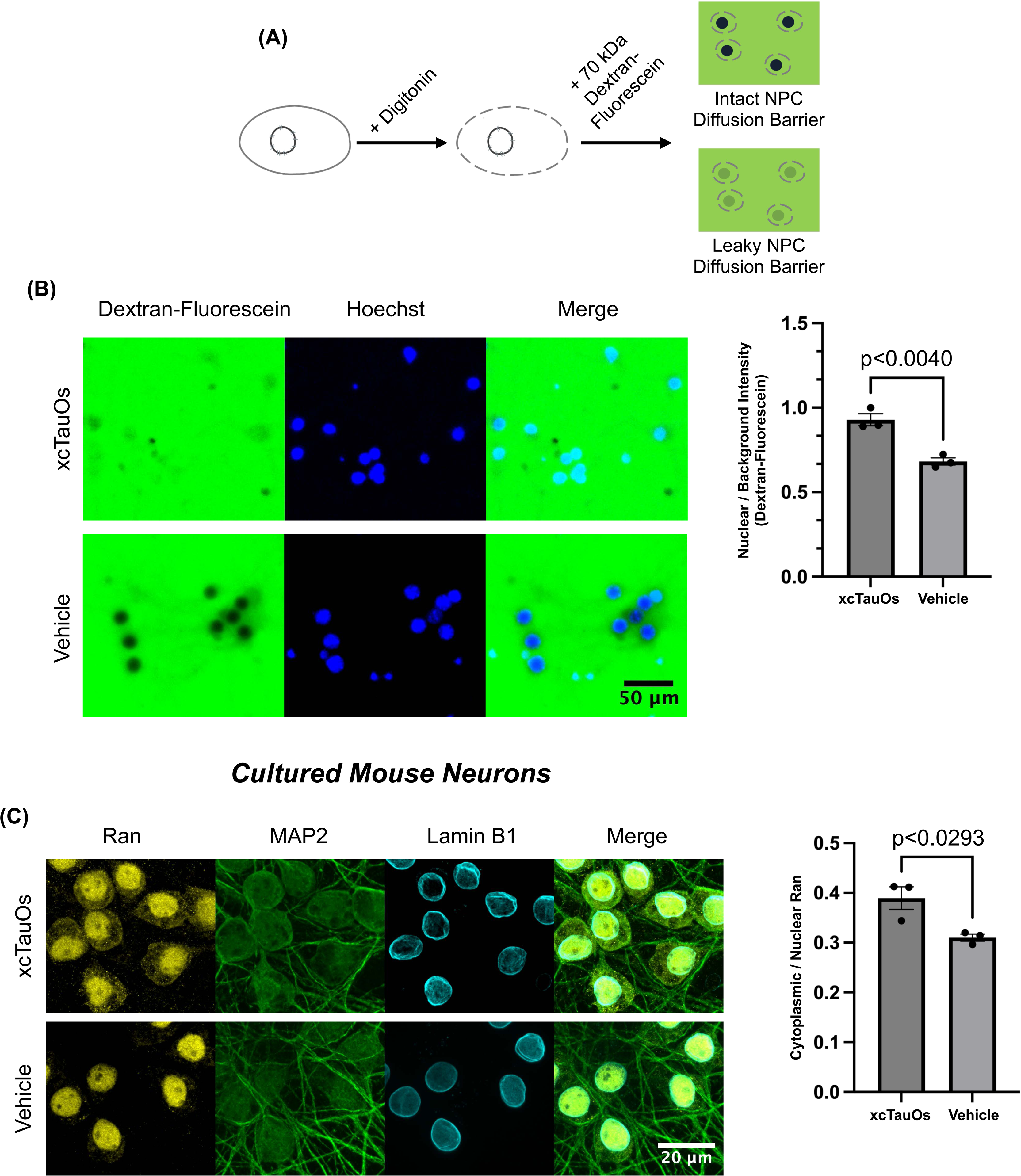
xcTauOs disrupt nucleocytoplasmic transport in primary WT mouse cortical neurons. (A) Schematic illustration of 70 kDa dextran-fluorescein exclusion assay. (B) Neurons treated with xcTauOs (250 nM total 2N4R tau) or vehicle for 24 hours had leaky NPCs. (C) Ran mislocalization caused by xcTauOs. Cultures were treated with xcTauOs (250 nM total 2N4R tau) or vehicle for 24 hours. Ran mislocalization was measured as the ratio of the mean pixel intensity of the cytoplasmic/nuclear Ran signal. Significance was determined using Student’s t test; error bars represent SEM; n = 3 biological replicates for A and B.

Next, we assessed the effect of xcTauOs on active nucleocytoplasmic transport in cultured mouse neurons by measuring the cytoplasmic/nuclear distribution of Ran, a small GTPase that reversibly shuttles between the nucleus and cytoplasm, but is normally highly enriched in nuclei at steady state [28, 29]. As shown in Figure 5C, xcTauOs caused an ∼26% increase in the cytoplasmic/nuclear Ran ratio relative to a vehicle control. Together, these dextran-fluorescein and Ran results demonstrate that xcTauOs functionally impair nucleocytoplasmic transport in neurons.

### 3.5 xcTauOs alter histone methylation and gene expression

Because the nuclear lamina serves as a tether for heterochromatin, we hypothesized that alterations of chromatin structure and gene expression accompanies the morphological distortion of neuronal nuclei caused by xcTauOs. To test that hypothesis, we first evaluated whether xcTauOs alter cultured mouse neuron levels of trimethylated lysine 9 of histone H3 (H3K9me3), which promotes transcriptional silencing. Supplementary Figure 5 shows that on western blots, the anti-H3K9me3 antibody used for these experiments was immunoreactive with bulk cell-derived histones, which include H3K9me3, but not with recombinant histone H3, which lacks post-translational modifications. This antibody also revealed by western blotting that the H3K9me3/total histone 3 ratio increased by ∼52% after xcTauO exposure (Figure 6A).

**FIGURE 6.**
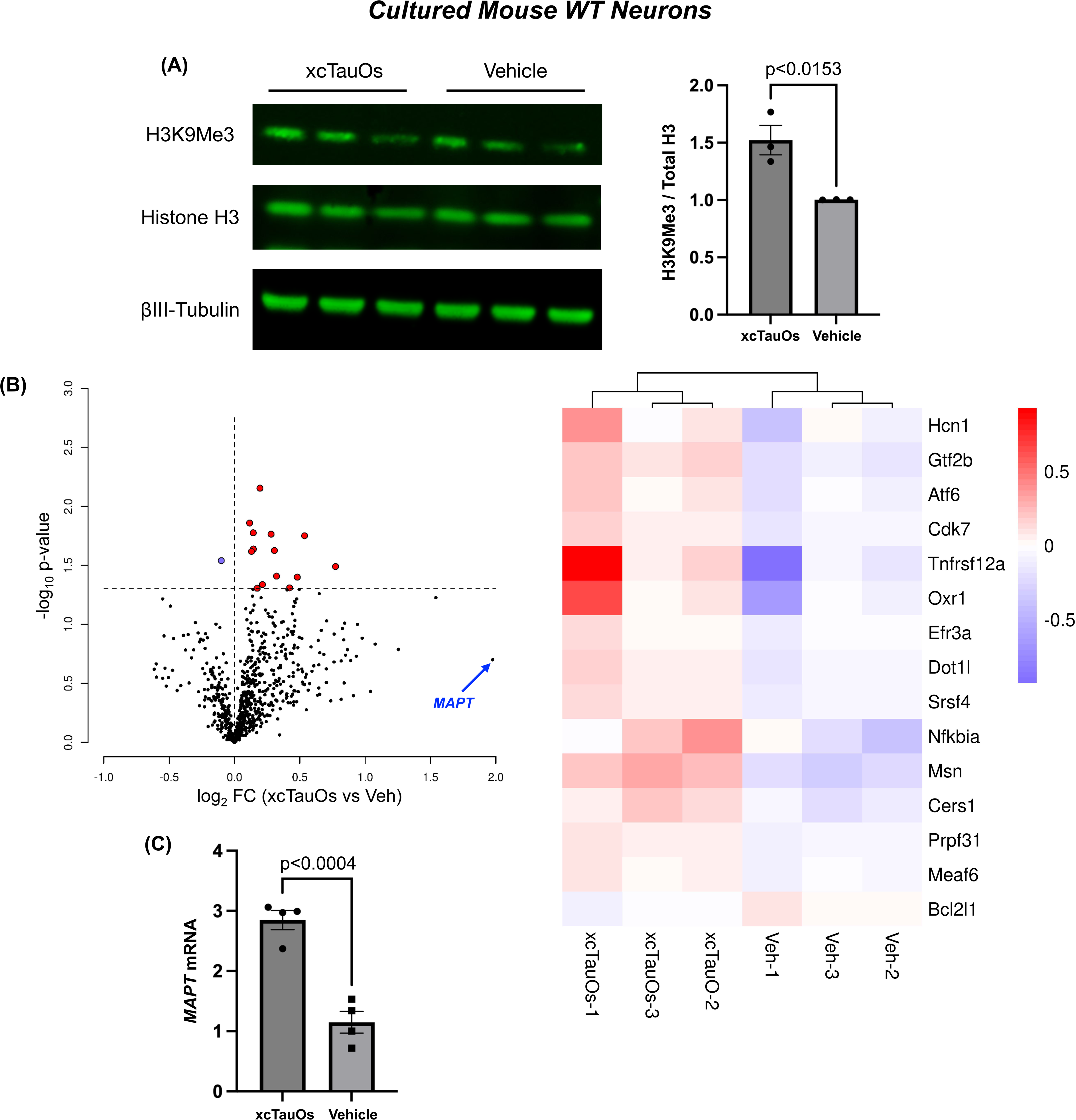
xcTauOs alter gene expression in primary mouse cortical neurons. (A) Western blot of H3K9Me3 for cultures treated xcTauOs (250 nM total 2N4R tau) or vehicle for 24 hours. Significance was determined using unpaired t test; error bars represent SEM; n = 3 biological replicates. (B) Volcano plot and heat map showing expression level changes detected by nanoSTRING nCounter technology for a panel of probes for 760 genes implicated in neurodegeneration. Cells were exposed to xcTauOs (250 nM total 2N4R tau) or vehicle for 6 hours; n = 3 biological replicates. (C) Quantification by qRT-PCR of *MAPT* mRNA levels in cells treated with xcTauOs (250 nM total 2N4R tau) or vehicle for 6 hours. Significance was determined using unpaired t test; error bars represent SEM; n = 4 biological replicates.

Next, we investigated whether xcTauOs alter gene expression in cultured neurons using the nCounter platform from nanoString for direct, unamplified measurement of 760 mRNAs associated with neuropathology. We found that for WT neurons, xcTauOs significantly altered mRNA levels for 15 genes, most of which were upregulated (Figure 6B). Annotation analysis of the differentially expressed genes highlights transcription functions based on genes like *Gtf2b*, *Srsf4*, *Prpf31* and *Cdk7*. In contrast, treatment of cultured tau^-/-^ neurons with xcTauOs yielded a different pattern: more genes were down- regulated than up-regulated, and none of genes affected in WT neurons were affected in tau^-/-^ neurons (Supplementary Figure 6). Interestingly, the most up-regulated gene in WT neurons after xcTauO treatment was *MAPT*, which encodes tau (fold change = 3.93; Figure 6B). However, *MAPT* elevation did not reach the significance threshold (p < 0.20), probably because of the high variation observed in this analysis. We therefore used an alternate method, qRT-PCR, to evaluate *MAPT* mRNA levels in WT neurons. That approach detected a statistically significant 2.84-fold increase of *MAPT* mRNA after xcTauO treatment of WT neurons (Figure 6C).

## 4 DISCUSSION

It is well established that tau can be released into the extracellular space by neurons independently of cell death and coupled to neuronal activity [30]. Much effort has focused on mechanisms of pathological tau transfer from neuron to neuron, but few cell biological responses of neurons to toxic tau uptake have been reported. We now provide evidence that xcTauOs provoke rapid structural and functional changes in neuronal nuclei. Within 1 hour of cultured neuron exposure to xcTauOs, their nuclei invaginate (Figures 2-4), and by no later than 24 hours of exposure, nucleocytoplasmic transport is impaired (Figure 5), and H3K9Me3 levels rise, and gene expression changes are evident (Figure 6). Intriguingly, the most prominent gene expression change is a nearly 3-fold increase in tau mRNA levels, which might signal a positive feedback loop whereby toxic TauOs drive their own expansion fueled not only by prion-like propagation [6–8], but by a growing pool of tau monomers as well. The likely *in vivo* significance of these results for cultured neurons is emphasized by our finding that invaginated neuronal nuclei are increased in human AD brain (Figure 1), consistent with a prior report [11], and in mouse models of AD (Supplementary Figure 2) and a pure tauopathy (Supplementary Figure 1). Altogether, these results implicate xcTauOs in the rapid transformation of healthy neurons into diseased neurons by changing nuclear architecture, and by extension, nuclear function.

Various forms of oligomerized tau were found to induce nuclear invagination in cultured neurons. Oligomers made from recombinant human 2N4R tau or an equimolar mix of all 6 human CNS tau isoforms were equally effective at 250 nM total tau (Figure 2C, D). Because only ∼1/3 of the total tau was typically oligomerized (Figure 2B), however, with dimers suspected of being most prevalent, the effective concentration of active oligomers was probably in the range of 30-40 nM. It is also noteworthy that TauOs made from all 6 human tau isoforms in hTau mice potently induced nuclear invagination (Figure 3A). This robust activity of TauOs made *in vivo* validates our use of TauOs made from recombinant human 2N4R tau for most experiments, especially since the activity of 2N4R TauOs was indistinguishable from those made from a mixture of all 6 recombinant human CNS tau isoforms.

Mechanistic insight into how xcTauOs cause nuclear invagination was provided by our finding that intracellular tau is required (Figure 4). In light of tau’s known prion-like behavior [6–8], it is significant that xcTauOs induced formation of perinuclear aggregates of endogenous tau in cultured mouse neurons (Figure 3C). Detailed structural analysis of these tau aggregates, and their potential roles in deforming and functionally compromising neuronal nuclei awaits further study.

A peak level of nuclear deformation in cultured neurons was reached within 1 hour of initial exposure to xcTauOs and was stable for at least 24 hours (Supplementary Figure 4). By that time point, and perhaps even earlier, a constellation of cell biological effects beyond nuclear deformation were evident. One example is impaired nucleocytoplasmic transport (Figure 5), which occurs at the nuclear pore complex (NPC). That finding is consistent with earlier reports of tau tightly associated with the NPC in frontotemporal dementia [9, 28] and directly interacting with Nup98, an NPC structural protein, to cause defective nucleocytoplasmic transport [9, 28]. This collection of results from our labs and others has profound implications. For example, multiple transcription factors, including p53, mislocalize from the nucleus to the cytoplasm in AD [31], perhaps as a consequence of nucleocytoplasmic transport disruption by xcTauOs.

We also found that xcTauOs increase the level of H3K9Me3, which epigenetically drives heterochromatin condensation and is elevated in AD brain [32]. Given the role of H3K9Me3 in controlling chromatin organization, it is not surprising that we observed multiple changes in gene expression following cultured neuron exposure to xcTauOs (Figure 6). We first explored fluctuations in mRNAs levels using the nanoString nCounter platform with a neuropathology panel that directly quantifies mRNAs encoded by 760 genes. Differential gene expression analysis after 6 hours of xcTauO exposure revealed 14 significantly up-regulated mRNAs and 1 that was significantly downregulated. Several of those up-regulated genes, like general transcription factor IIB (Gtf2b), Ser/Arg-splicing factor 4 (Srsaf4) and cyclin dependent kinase 7 (Cdk7), are associated with transcription and splicing annotations. Consistent with our finding that nuclear invagination by xcTauOs requires intracellular tau (Figure 4), none of the genes whose transcription levels were affected by xcTauOs in WT neurons was altered in tau^-/-^ neurons (Supplementary Figure 6).

Remarkably, the most up-regulated gene in WT neurons following xcTauO treatment was *MAPT*, which encodes tau itself. Its upregulation did not reach statistical significance for the nCounter experiments, though, because one of the 3 biological replicates yielded an outlier result. Accordingly, we also used qRT-PCR to measure tau mRNA in 4 biological replicates, all of which were distinct from the samples analyzed using nCounter. That approach indicated a statistically significant 3-fold increase in *MAPT* mRNA levels as a consequence of xcTauO treatment of WT neurons. When considered together, the nCounter and qRT-PCR results point to the likelihood that xcTauOs induce a large increase in mRNA for tau. In such a scenario, xcTauOs might trigger a positive feedback loop that stimulates production of excess tau mRNA, and by extension, more tau protein and toxic TauOs.

## ACKNOWLEDGEMENTS

Funding for this work was provided by NIH Grant AG051085 (GSB), the Owens Family Foundation (GSB), the Cure Alzheimer’s Fund (GSB), the Rick Sharp Alzheimer’s Foundation (GSB), Webb and Tate Wilson, and the NanoString nCounter Grant Program for the University of Virginia’s Spatial Biology Core. We would like to thank Dr. Dora Bigler-Wang for handling mice and preparing primary neuron cultures; Dr. Heather Ferris for providing human brain tissues for western blots; Dr. Anthony Spano for providing βIII-tubulin antibody (Tuj1); the late Dr. Lester (Skip) Binder for providing us Tau5 hybridoma cells; Drs. Michael Vitek and Carol Colton for their prior gift of CVN mice; and Drs. John S. Lazo and Elizabeth R. Sharlow for their invaluable intellectual involvement throughout this project. This paper partially fulfills the Ph.D. requirements for Xuehan Sun, and we thank all members of her thesis dissertation committee not mentioned otherwise here for their conscientious and wise counsel: Drs. Christopher Deppmann, Xiaorong Liu and Thurl Harris.

## AUTHOR CONTRIBUTIONS

XS co-conceived, co-designed and performed the bulk of the research, and co-analyzed the data. YS and SS performed and co-analyzed the immunohistochemistry data. GE and AK performed and co-analyzed the nCounter data. JWM provided annotated human brain samples for immunohistochemistry and assisted with interpretation of the corresponding micrographs. JRL provided PS19 brain sections, and along with AN, provided ongoing counsel about experimental design and data interpretation. GSB co-conceived, co-designed and supervised the research. XS, GE, and GSB wrote the manuscript. All authors participated in editing the manuscript and approved the submitted version.

## SUPPLEMENTARY MATERIALS

**SUPPLEMENTARY FIGURE 1.**
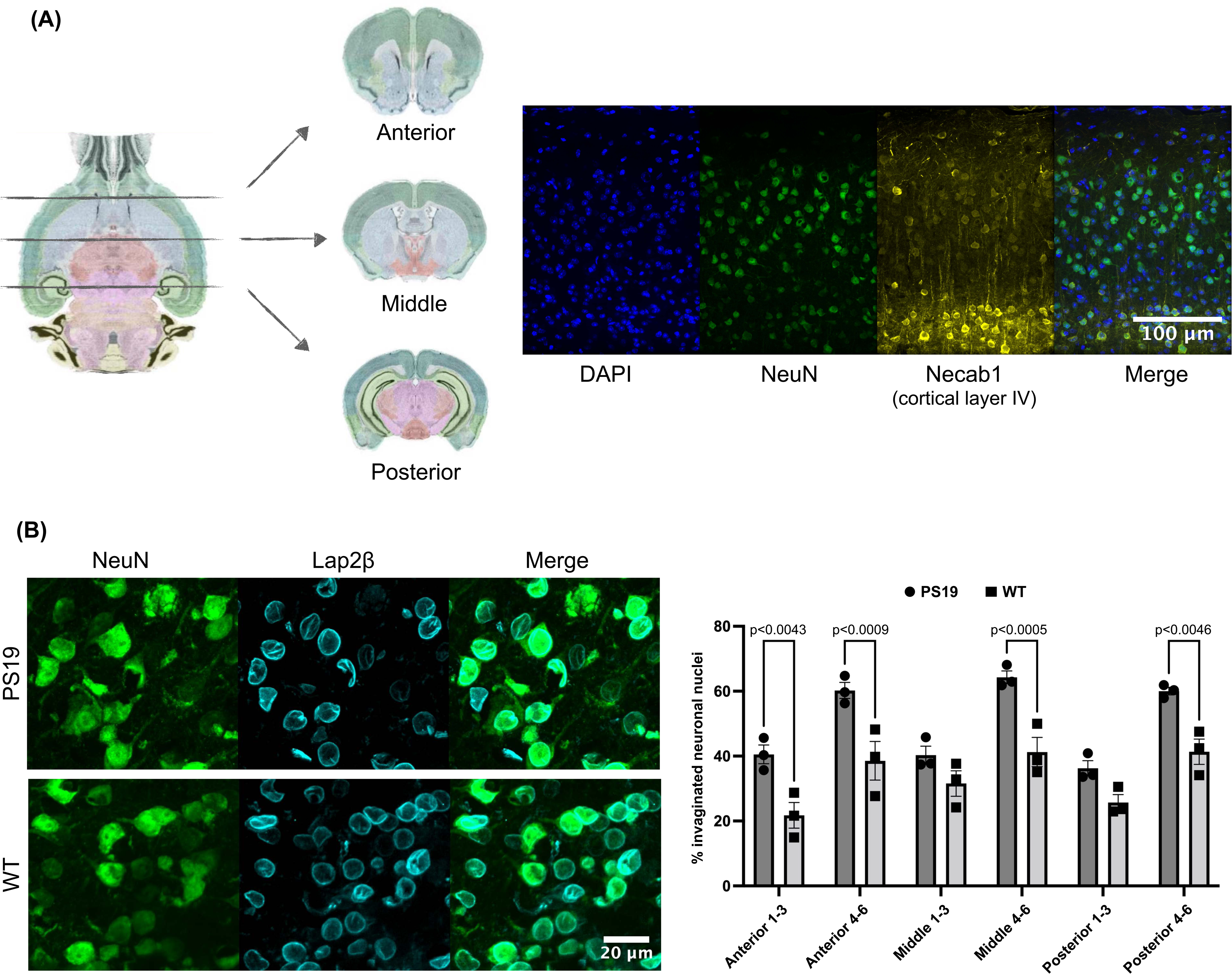
Neuronal nuclear invagination in 6-month-old PS19 mouse brain. (A) Schematic of sample collection (image credit: Allen Institute for Brain Science https://mouse.brain-map.org/agea), and immunostaining for Necab1, a marker of cortical layer IV neurons, and for the neuronal nucleus marker, NeuN. (B) Incidence of neuronal nuclear invaginations in deep cortical layers. Significance was determined using two-way ANOVA followed by Bonferroni’s test; error bars represent SEM; n = 3 biological replicates, and about 300 neurons from each section were evaluated.

**SUPPLEMENTARY FIGURE 2.**
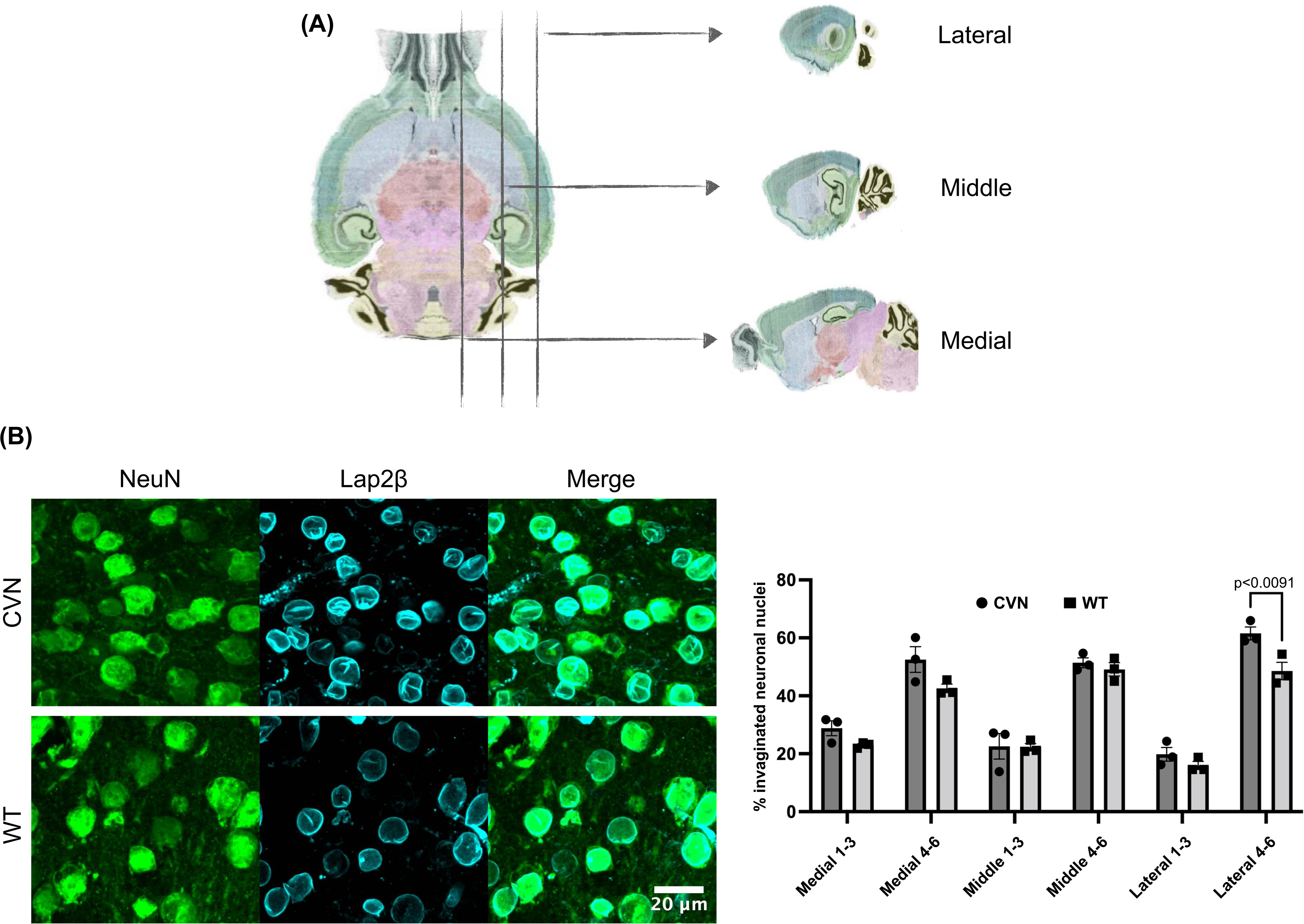
Neuronal nuclear invagination in 22-month-old CVN mouse brain. (A) Schematic of sample collection (image credit: Allen Institute for Brain Science https://mouse.brain-map.org/agea). (B) Incidence of neuronal nuclear invaginations in lateral deep cortical layers. Significance was determined using two-way ANOVA followed by Bonferroni’s test; error bars represent SEM; n = 3 biological replicates, and about 300 neurons from each section were evaluated.

**SUPPLEMENTARY FIGURE 3.**
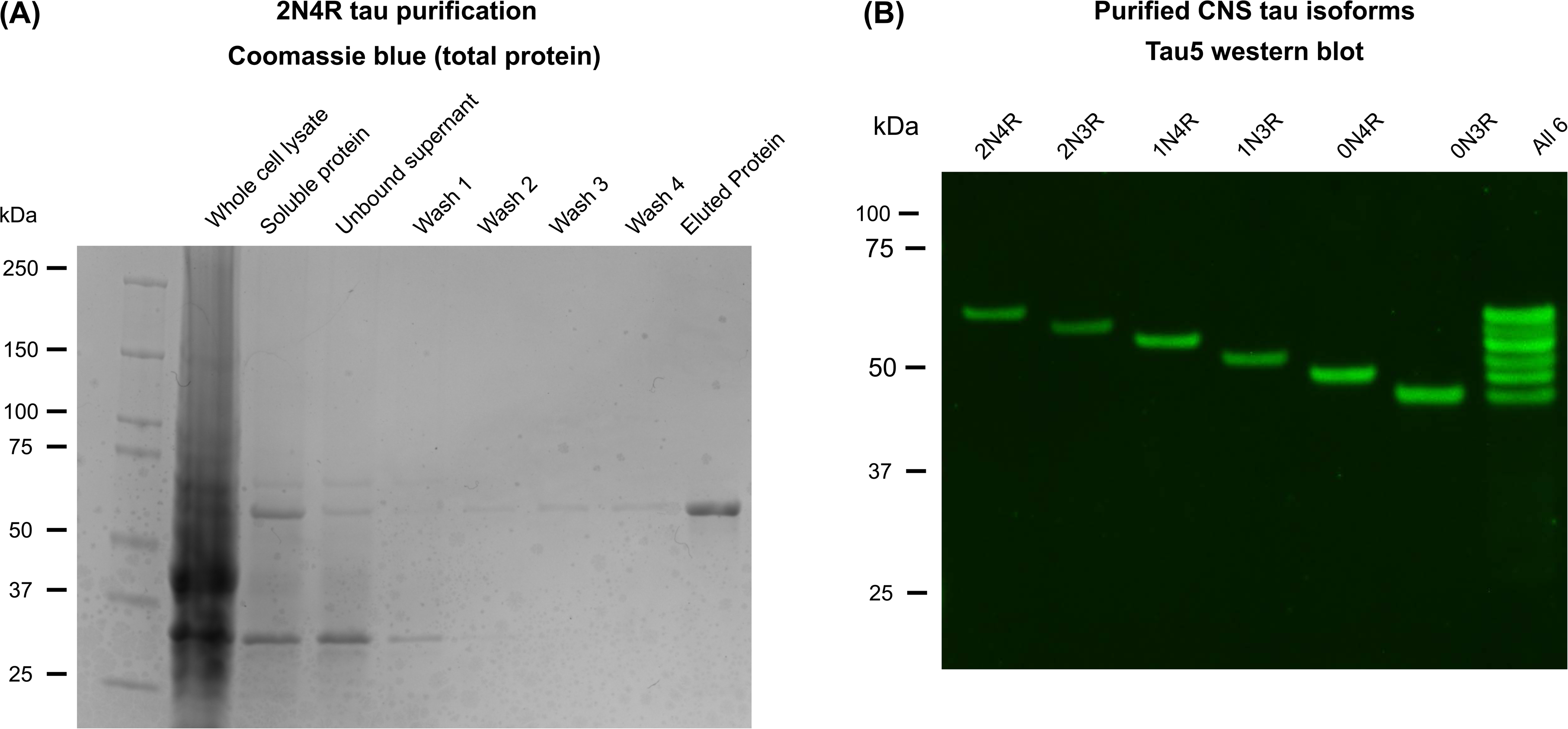
Recombinant his-tagged human tau used in this study. (A) 2N4R tau purification using Talon metal affinity chromatography. (B) Tau5 western blot of purified versions of each of the 6 human CNS tau isoforms.

**SUPPLEMENTARY FIGURE 4.**
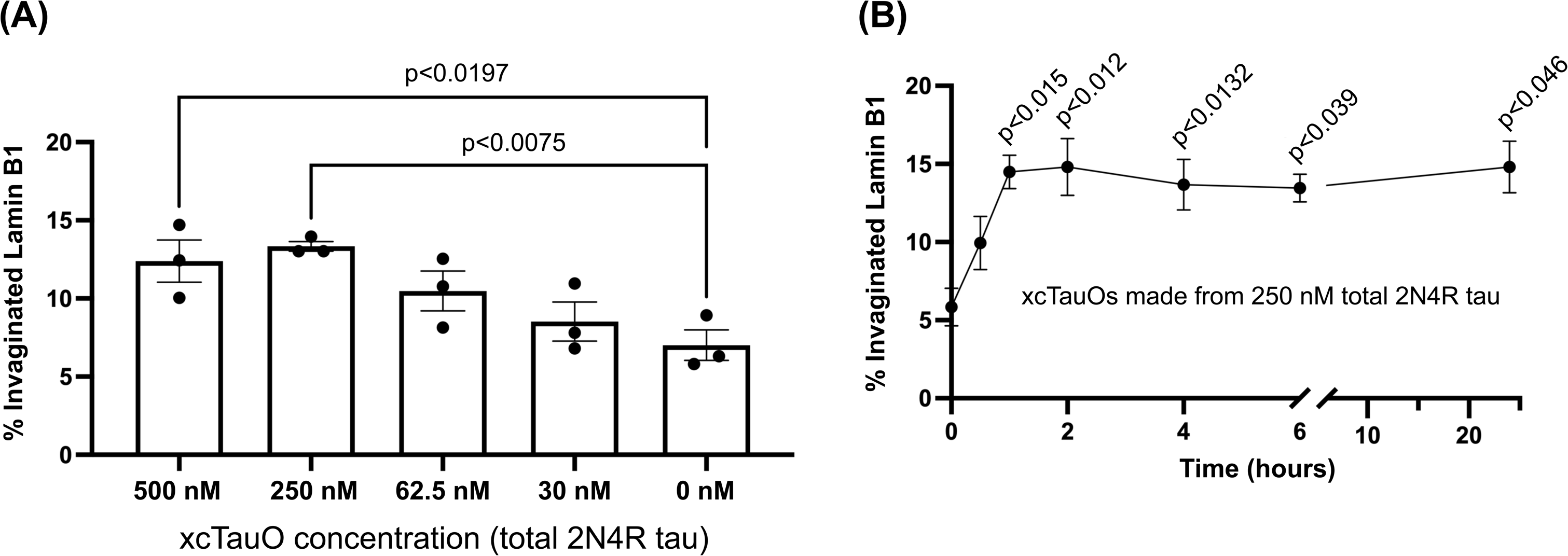
Dose response and kinetics of nuclear invagination in primary WT mouse cortical neurons caused by xcTauOs (2. (A) Dose response for a 1-hour exposure. (B) Kinetics of nuclear invagination caused by xcTauOs (250 nM total 2N4R tau). Significance was determined using one-way ANOVA followed by Dunnett’s test; error bars represent SEM; n = 3 independent experiments for A and B.

**SUPPLEMENTARY FIGURE 5.**
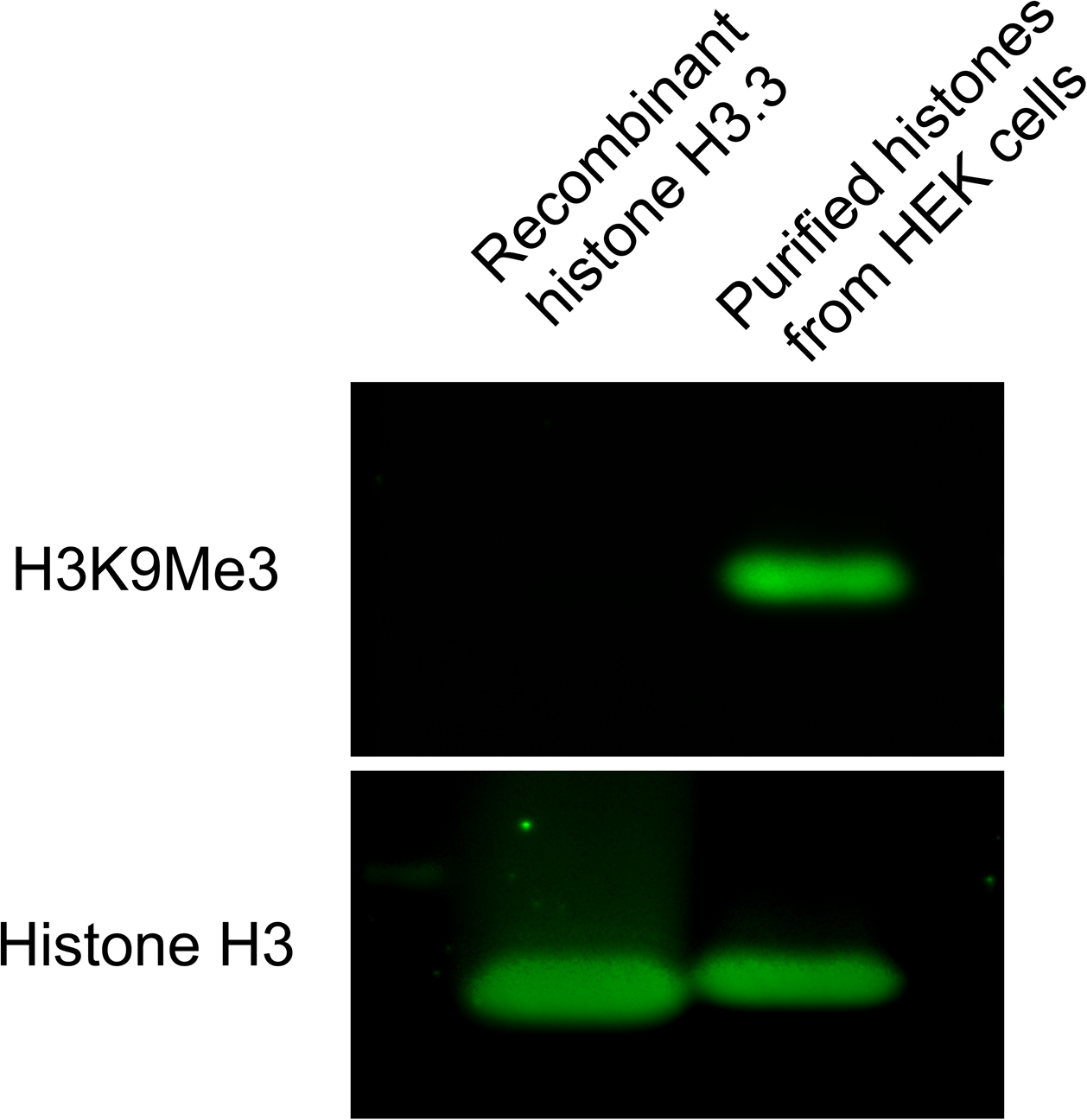
Validation of H3K9Me3 antibody. Western blot analysis of unmodified recombinant histone and native histones (bearing a full repertoire of post-translational modifications) extracted from HEK cells.

**SUPPLEMENTARY FIGURE 6.**
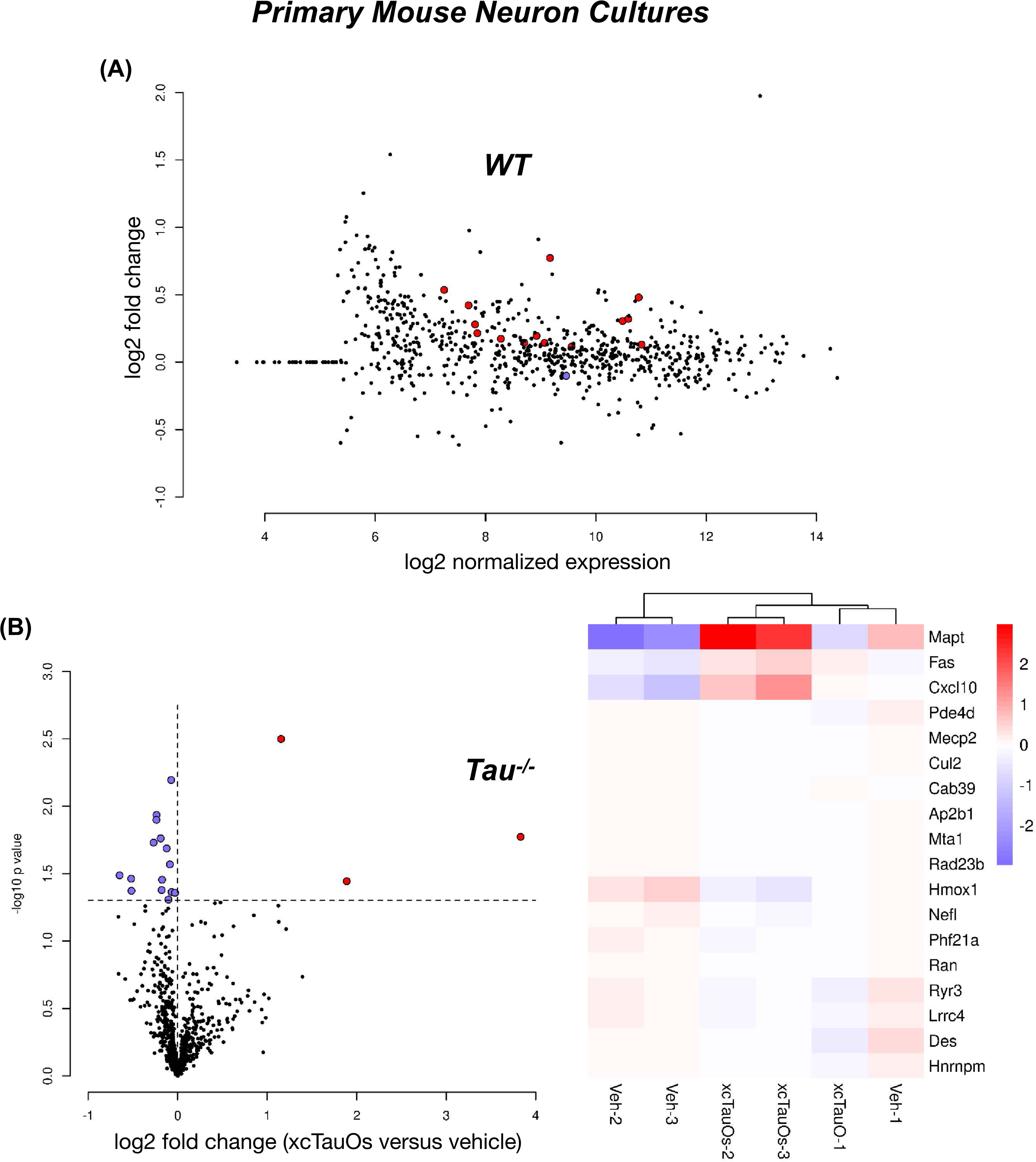
Differential gene expression caused by xcTauOs in WT and tau^-/-^ neurons detected by nanoSTRING nCounter analysis of 760 genes implicated in neurodegeneration. (A) The relationship between expression and fold-change for WT neurons exposed for 6 hours to xcTauOs (250 nM total 2N4R tau) versus vehicle. (B) Volcano plot comparing of tau^-/-^ neurons treated for 6 hours with xcTauOs (250 nM total 2N4R tau) or vehicle, and corresponding heat map.

**SUPPLEMENTARY TABLE 1.**
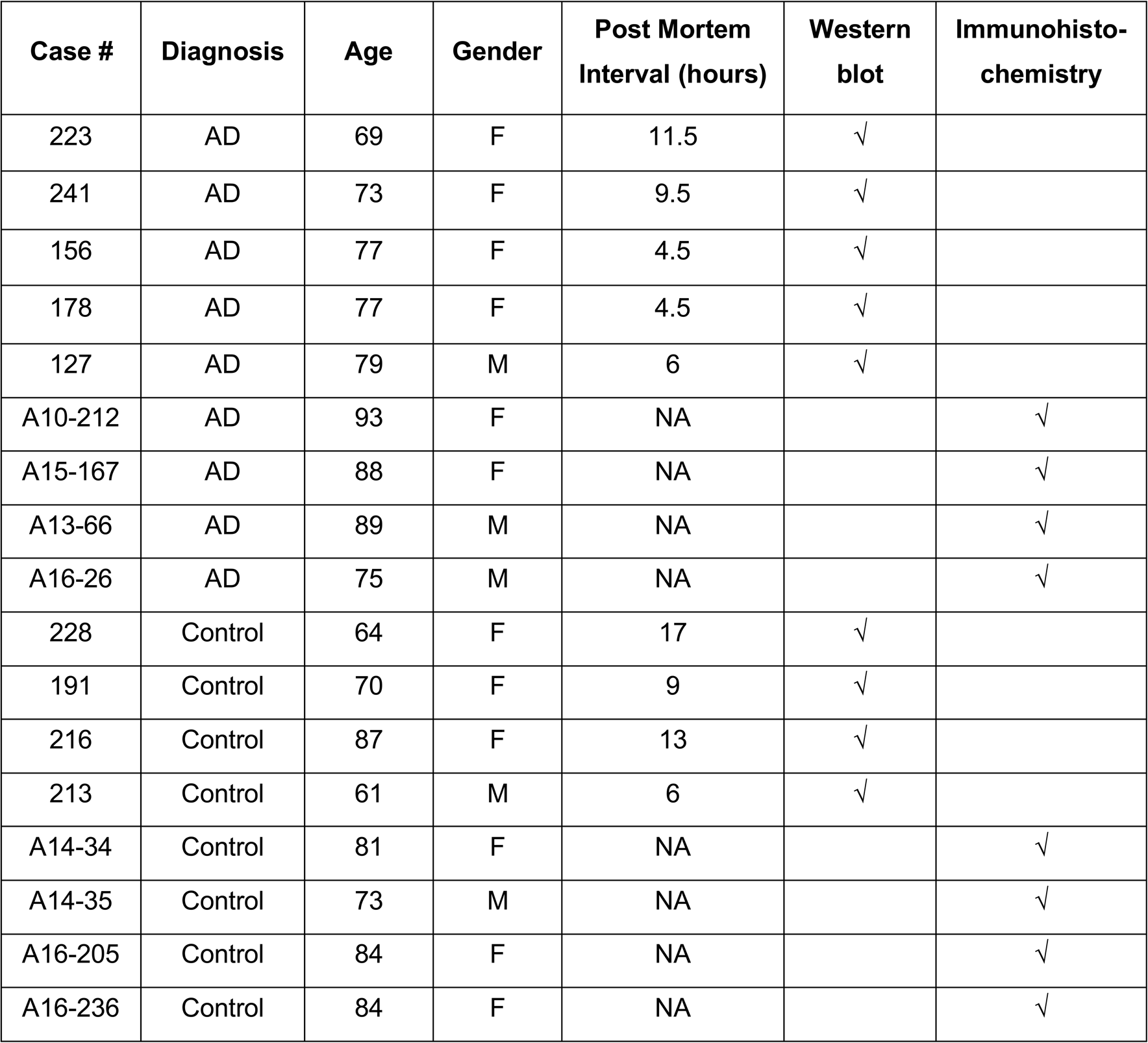
Patient demographics and tissue-specific applications.

| Case # | Diagnosis | Age | Gender | Post Mortem Interval (hours) | Western blot | Immunohisto-chemistry |
| --- | --- | --- | --- | --- | --- | --- |
| 223 | AD | 69 | F | 11.5 | √ |  |
| 241 | AD | 73 | F | 9.5 | √ |  |
| 156 | AD | 77 | F | 4.5 | √ |  |
| 178 | AD | 77 | F | 4.5 | √ |  |
| 127 | AD | 79 | M | 6 | √ |  |
| A10-212 | AD | 93 | F | NA |  | √ |
| A15-167 | AD | 88 | F | NA |  | √ |
| A13-66 | AD | 89 | M | NA |  | √ |
| A16-26 | AD | 75 | M | NA |  | √ |
| 228 | Control | 64 | F | 17 | √ |  |
| 191 | Control | 70 | F | 9 | √ |  |
| 216 | Control | 87 | F | 13 | √ |  |
| 213 | Control | 61 | M | 6 | √ |  |
| A14-34 | Control | 81 | F | NA |  | √ |
| A14-35 | Control | 73 | M | NA |  | √ |
| A16-205 | Control | 84 | F | NA |  | √ |
| A16-236 | Control | 84 | F | NA |  | √ |

**SUPPLEMENTARY TABLE 2.** Primary and secondary antibodies.

| Name | Tag | Source;Cat # |
| --- | --- | --- |
| chicken anti-MAP2 | None | abcam; ab92434 |
| Mouse anti-pan-tau (Tau5) | None | Dr. Lester (Skip) Binder (deceased) |
| mouse anti-NeuN | None | Millipore; MAB377 |
| mouse anti-βIII-tubulin | None | Dr. Tony Spano |
| mouse anti-H3K9me3 | None | Novus Biologicals; NBP1-30141 |
| mouse anti-Ran | None | BD Biosciences; 610341 |
| mouse anti-Lap2β | None | BD Biosciences; 611000 |
| Mouse anti-tau (T49) | None | Millipore; MABN827 |
| rabbit anti-LaminB1 | None | abcam; ab16048 |
| rabbit anti-H3 | None | Cell Signaling Technology; #9715 |
| rabbit anti-Necab1 | None | Sigma-Aldrich; HPA023629 |
| rabbit anti-HSP90 | None | Cell Signaling Technology; C45G5 |
| Goat anti-Mouse IgG | Alexa Fluor 488 | Invitrogen; A-11001 |
| Goat anti-Chicken IgY | Alexa Fluor 568 | Invitrogen; A-11041 |
| Goat anti-Rabbit IgG | Alexa Fluor 647 | Invitrogen; A-21244 |
| Goat anti-Rabbit IgG | IRDye 680RD | LI-COR; 926-68071 |
| Goat anti-Mouse IgG | IRDye 800CW | LI-COR; 926-32210 |

